# FANCD2-FANCI is a clamp stabilized on DNA by monoubiquitination during DNA repair

**DOI:** 10.1101/858225

**Authors:** Pablo Alcón, Shabih Shakeel, Ketan J. Patel, Lori A. Passmore

**Author notes:** These authors contributed equally (Pablo Alcón, Shabih Shakeel).

## Abstract

Vertebrate DNA crosslink repair is a crucial process that excises toxic replication-blocking DNA interstrand crosslinks^1^. This pathway fails in the inherited human disease Fanconi anemia (FA), resulting in abnormal development, loss of blood production, and marked cancer susceptibility^2^. Numerous factors involved in crosslink repair have been identified, and mutations in their corresponding genes cause FA^3^. A biochemical description of how FA gene products might initiate the removal of DNA crosslinks is now emerging. However, structural insight into this vital process has been limited. Here, we use electron cryomicroscopy (cryoEM) to determine the structure of a key DNA crosslink repair factor – FANCD2 heterodimerized with FANCI (D2-I). Recombinant chicken D2-I adopts a conformation that is in agreement with the structure of its murine counterpart^4^. In contrast, the activated monoubiquitinated form of D2-I (ubD2-I) adopts an alternative closed conformation, creating a channel that encloses double-stranded DNA. Ubiquitin is positioned at the interface of FANCD2 and FANCI, and acts as a molecular pin to trap ubD2-I in the closed conformation, clamped on DNA. The new solvent-exposed interface around the monoubiquitination site is likely a platform for recruitment of other DNA repair factors. We also find that isolated FANCD2 is a dimer, but it adopts a closed conformation that is unable to bind DNA. When incubated with free FANCI, the FANCD2 homodimer exchanges to D2-I, acquiring DNA binding properties. This suggests an autoinhibitory mechanism that prevents FANCD2 activation on DNA until after assembly with FANCI. Together, our cryoEM and biochemical analysis suggests that D2-I is a clamp that is locked onto DNA by ubiquitin, and provides unanticipated new insight into the regulation and the initiation of DNA cross-link repair.

---

A key step in the FA DNA crosslink repair pathway is the monoubiquitination of the FANCD2-FANCI heterodimer^5-9^. Monoubiquitination occurs during DNA replication and is stimulated by the FA core complex – a multi-subunit DNA damage inducible monoubiquitin ligase. Activated ubD2-I localizes to sites of DNA damage and recruits nucleases to remove the lesion^10-17^. Monoubiquitination is tightly controlled, and can be recapitulated in cell free systems in the presence of DNA^18,19^. ubD2-I must be deubiquitinated by USP1/UAF1 for completion of crosslink repair^20-22^. A longstanding mystery is why D2-I monoubiquitination and its subsequent reversal should be so fundamental for crosslink repair. Moreover, the mechanistic details of FANCD2 and FANCI interaction with DNA, and how this might ultimately enable DNA crosslink removal remain unclear. Here, we have expressed, purified, and monoubiquitinated *Gallus gallus* FANCD2 and D2-I complexes, and, using cryoEM and biochemical assays, we delineated key details of how these critical factors interact with DNA.

First, we purified a recombinant D2-I complex after co-expression of both proteins in insect cells, and incubated it with His-tagged ubiquitin, E1, E2 (UBE2T), recombinant FA core complex^23^ and DNA (Fig. 1a; Extended Data Fig. 1a). This resulted in specific monoubiquitination of FANCD2 on K563^23^. We enriched for ubD2-I by affinity chromatography using the His-tag on ubiquitin (Fig. 1b). Next, we used cryoEM to investigate the architecture of both unmodified D2-I and activated ubD2-I. We obtained 3D structures of both complexes at overall resolutions of ∼4 Å (Fig. 1c, d; Extended Data Fig. 1b-e; Extended Data Table 1). The structure of unmodified D2-I agrees with the previously reported crystal structure of the *Mus musculus* proteins^4^. FANCD2 and FANCI share a striking structural similarity containing a series of α-helices arranged into four superhelical α-solenoid structures (S1-S4) and two helical domains (HD1, HD2). They fold into an overall shape akin to two antiparallel saxophones. In this arrangement, the ubiquitination sites (D2_K563_ and I_K525_ for the chicken proteins) are buried within the dimerization interface, indicating that remodeling of the complex is very likely required for access by the FA monoubiquitin ligase complex.

**Fig. 1:**
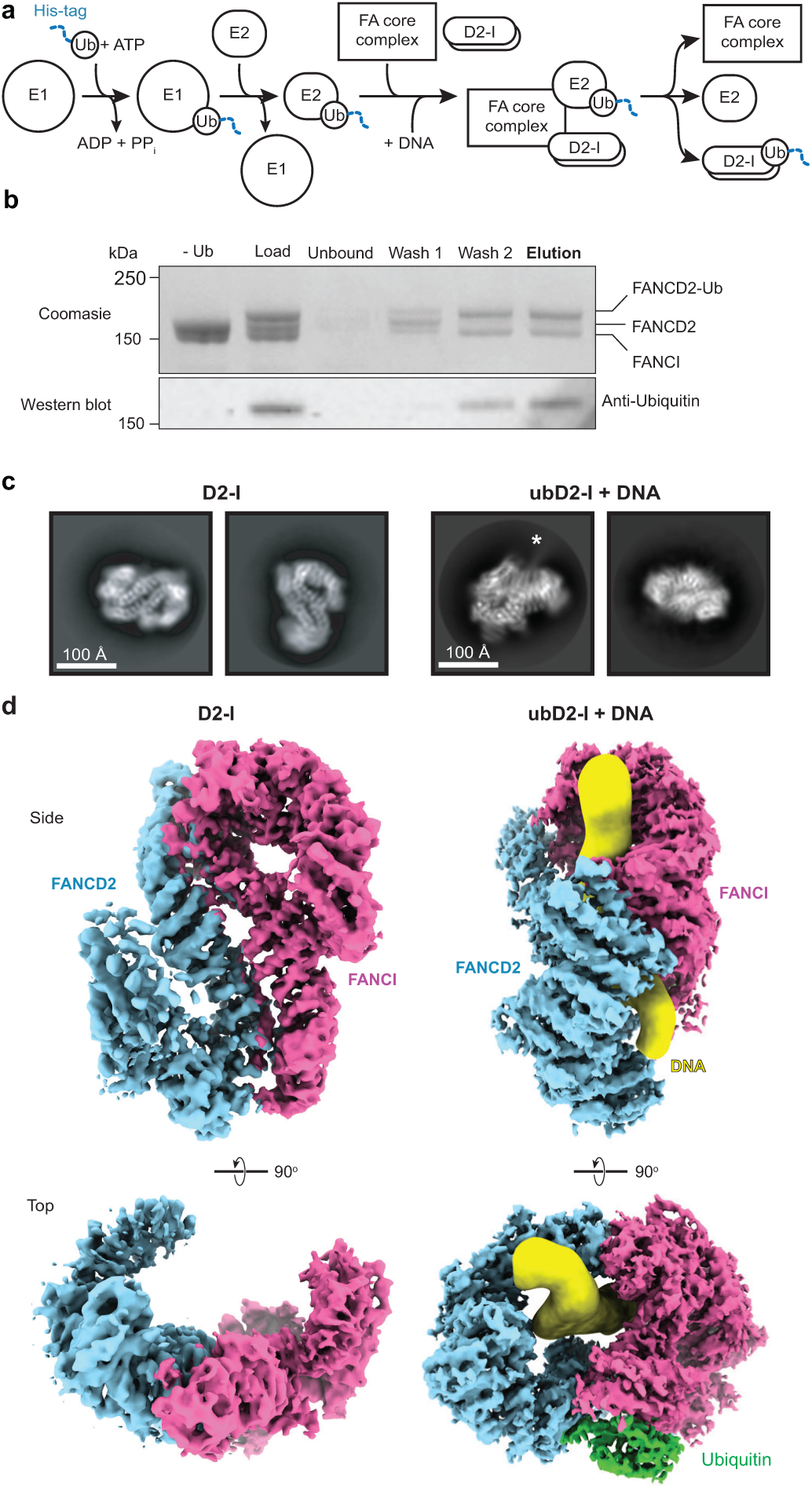
Purification and structure of D2-I and ubD2-I. **a** Scheme for monoubiquitination of D2-I using fully recombinant components. The His-tag on ubiquitin is shown as a blue dashed line. **b** ubD2-I was enriched by purification of His-tagged ubiquitin on Ni-NTA. The unbound fraction, two wash fractions and elution were analyzed by Coomassie-stained SDS-PAGE (top) and Western blotting with an anti-Ubiquitin antibody (bottom). **c** Selected 2D reference-free class averages of D2-I (left) and ubD2-I with DNA (right). An asterisk marks density extending from the side of the complex that we assign to DNA. **d** CryoEM maps of D2-I (left) and ubD2-I with DNA (right), segmented into FANCD2 (blue), FANCI (magenta), ubiquitin (green) and DNA (yellow).

In striking contrast, the structure of ubD2-I is markedly different from the unmodified complex. Symmetry was not obvious in 2D class averages (Fig. 1c), but the 3D reconstruction showed that the overall ubD2-I complex retains some pseudo-symmetric features (Fig. 1d). In the monoubiquitinated form, FANCD2 and FANCI are folded towards each other so the C-terminal domains of each monomer (akin to the bells of the saxophone), wrap around density that we assigned to double-stranded DNA (see below). In support of this, multi-body refinement of ubD2-I showed that there is variability in the relative positions of FANCD2 and FANCI (Supplementary Video 1), and this is consistent with previous data^24^.

We then built homology models of FANCD2, FANCI and ubiquitin, and fitted these into the D2-I and ubD2-I maps (Fig. 2a; Extended Data Fig. 2-4). Superimposition of the two models showed that the dimer interface acts as a hinge, allowing the C-termini of FANCD2 and FANCI to swing towards each other, rotating by almost 70° (Supplementary Videos 2-4). This conformational change in ubD2-I exposes the K563 monoubiquitination site in FANCD2 on the backside of the hinge, where we unambiguously identified density for ubiquitin in the cryoEM map (Fig. 2b). Ubiquitin is covalently anchored onto FANCD2 (Fig. 1b) but it docks onto the surface of FANCI, and likely stabilizes the complex. In contrast, the FANCI monoubiquitination site (K525) is only partially exposed (Extended Data Fig. 2c), consistent with inefficient *in vitro* monoubiquitination of FANCI with recombinant or native FA core complex^18,23,25^ (Fig. 1b). In cells, FANCI monoubiquitination is stimulated by phosphorylation but is not essential for DNA crosslink repair^9,26^.

**Fig. 2:**
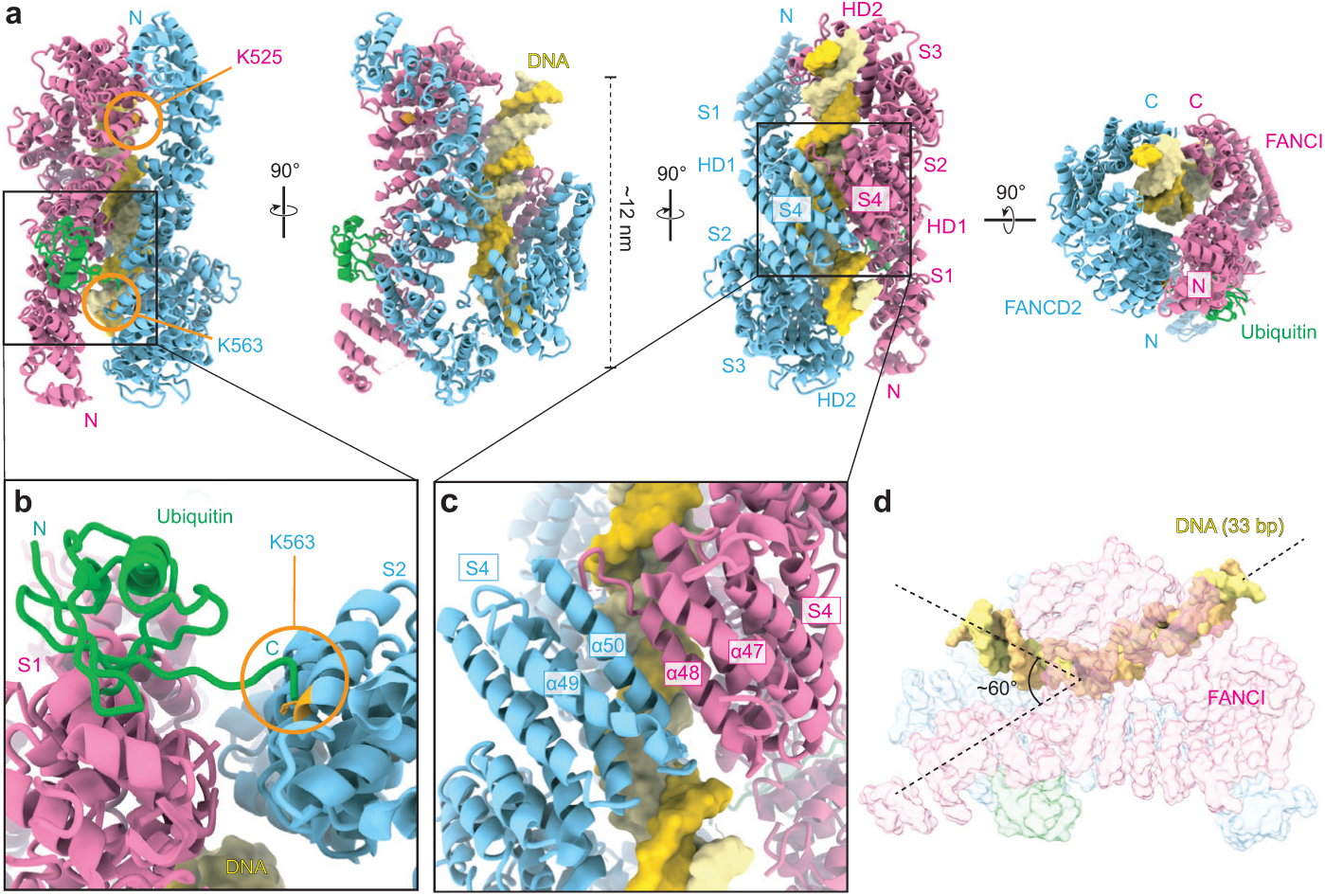
ubD2-I is a DNA clamp. **a** Homology models for FANCD2 (blue), FANCI (magenta) and ubiquitin (green) were fit into the ubD2-I cryoEM map. The N- and C-termini, solenoids 1-4 (S1-S4) and helical domains (HD1, HD2)^4^ of FANCD2 and FANCI are indicated. The monoubiquitinated lysine in FANCD2 and the lysine that can be monoubiquitinated in FANCI are shown in orange. A model of 33 bp double-stranded DNA (yellow) was also placed into the map. **b** Close-up view of the monoubiquitination site. The ubiquitin moiety attached to FANCD2 makes extensive contacts with FANCI. **c** Close-up view of the C-terminal domains (S4) of ubD2-I that clamp around DNA. **d** Double-stranded DNA is kinked within the ubD2-I complex.

The C-terminal solenoids (S4) of FANCD2 and FANCI, as well as additional unmodeled C-terminal regions, mediate contacts at the new interface (Fig. 2c). This interaction likely stabilizes the C-terminal region of FANCD2, which becomes well ordered in ubD2-I map (Fig. 1d). The repositioned C-terminal domains of FANCD2 and FANCI in the monoubiquitinated complex form a new channel between the two protomers. The channel contained density for ∼15-20 bp double-stranded DNA. The DNA extended beyond the channel and we could model a total of 33 bp. The DNA was at lower resolution than the rest of the map, likely because of heterogeneity in the DNA position, and inherent flexibility. DNA makes contacts with both proteins, but it appears to be shifted towards FANCI (Fig. 2a). Contacts are likely mediated largely through interactions between the negatively-charged phosphate backbone and positively-charged surface residues^4^. Interestingly, the bound DNA appears to be kinked next to the C-terminal domains within the ubD2-I complex (Fig. 2d).

DNA is required for monoubiquitination of D2-I *in vitro*^18,19^ but the role of DNA in this reaction was unclear. To further investigate this, we performed monoubiquitination assays of D2-I in the presence of a series of double-stranded DNAs of varying lengths. D2-I was monoubiquitinated in the presence of a 19 bp DNA, but not with a 14 bp DNA (Extended Data Fig. 5a). We also analyzed protein–DNA interactions in the monoubiquitination assays using electrophoretic mobility shift assays (EMSAs). D2-I did not bind to a 14 bp DNA, but interacted efficiently with DNA substrates that were 19 bp or longer (Extended Data Fig. 5a). Therefore, these biochemical data show that the minimum DNA length required for ubiquitination (15– 19 bp) correlates with the minimum length required for D2-I binding. Moreover, this agrees well with the length of DNA encircled by ubD2-I in the cryoEM structure, and with the distance between a stalled replication fork and a DNA crosslink^6^.

Together, these experiments explain why and how DNA stimulates FANCD2 monoubiquitination: D2-I binds double-stranded DNA, promoting closure of the heterodimer to expose the K563 monoubiquitination site in FANCD2 and allow access to the activated E2 enzyme. In addition, the structures and biochemical studies suggest that monoubiquitination of FANCD2 on the backside of the hinge acts as a covalent molecular pin, locking the closed heterodimer on DNA and preventing it from rotating back to the open D2-I conformation. In agreement with this, monoubiquitination results in a small, but reproducible, increase in DNA binding affinity (Extended Data Fig. 5b). To further test this hypothesis, we incubated D2-I with FAM-labeled DNA and then tested the ability of a second (Alexa-labeled) DNA to be incorporated into D2-I by displacing the FAM-DNA. We found that in the absence of ubiquitination, the second DNA is readily incorporated into D2-I. In contrast, when FAM-DNA is used to form the ubD2-I complex, a negligible amount of the competing DNA is incorporated (Extended data Fig. 5d).

A major remaining question was why FANCD2 was not monoubiquitinated *in vivo* or *in vitro* in the absence of FANCI^13,18,19^. To address this, we purified FANCD2 and FANCI separately. Purified FANCI ran as a monomer on gel filtration chromatography whereas, unexpectedly, FANCD2 ran as a dimer (Fig. 3a). Next, we assessed their DNA binding capacities and found that FANCI bound DNA efficiently but FANCD2 did not interact with DNA at any of the concentrations we tested (Fig. 3b; Extended Data Fig. 5c). Moreover, FANCD2 was not substantially monoubiquitinated by the FA core complex, and this could not be stimulated by DNA (Extended Data Fig. 6a). To understand the molecular basis of FANCD2 dimerization and gain insight into why it did not bind DNA, we determined the structure of the FANCD2 homodimer using cryoEM to ∼4 Å resolution (Fig. 3c; Extended Data Fig. 1). The FANCD2 dimer was symmetric with each protomer containing the same overall domain architecture as in the heterodimeric complex. Surprisingly, the FANCD2 homodimer was in a closed conformation, more similar to ubD2-I than to D2-I. This provides a molecular explanation for the lack of DNA binding by FANCD2: The DNA binding sites were blocked because it was in a closed conformation that presumably does not open to allow entry of a double-stranded helix. In addition, FANCD2 homo-dimerization occludes K563 on both protomers, potentially preventing their ubiquitination (Fig. 3c; Extended Data Fig. 5a).

**Fig. 3:**
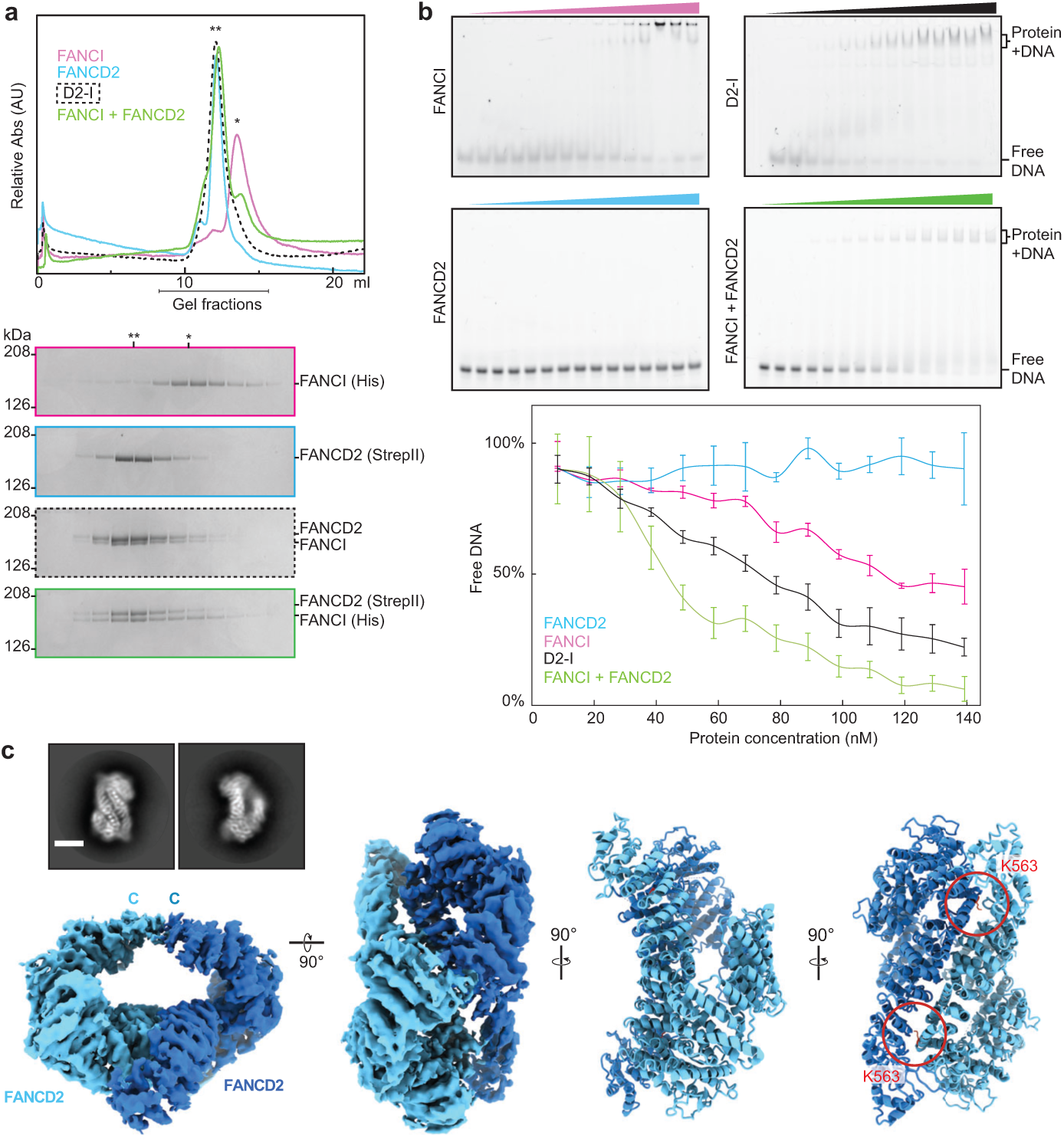
FANCI forms a monomer and binds DNA whereas FANCD2 is dimeric and does not bind DNA. **a** Analysis of purified FANCI, FANCD2, D2-I, and FANCI mixed with FANCD2 by size exclusion chromatography. Chromatographs (top) and SDS-PAGE analysis of fractions (bottom) are shown. A single asterisk (*) indicates the migration position for monomers. A double asterisk (**) indicates the migration position for dimers. This experiment was performed three times. **b** DNA binding of FANCI, FANCD2, D2-I, and FANCI mixed with FANCD2 was analyzed by EMSAs performed with 20 nM 39 bp double-stranded DNA and 0–140 nM protein. Representative gels of experiments independently performed three times (top) and quantitation of mean intensity of free DNA (bottom) are shown. Error bars represent the standard deviation. Data points are connected by lines for clarity. **c** CryoEM map and model of FANCD2 dimer. The location of the C-terminus is marked. Selected 2D reference-free class averages (inset) are also shown. The scale bar is 100 Å. The buried K563 on either FANCD2 protomer are shown in red.

The FANCD2 homodimer was stable over a gel filtration column, indicative of a high affinity interaction. To investigate how the heterodimeric D2-I complex forms, we mixed purified FANCD2 and FANCI and analyzed their oligomerization state. We found that after incubation with FANCD2, the migration position of FANCI on a gel filtration column shifted to the position of a dimer (Fig. 3a). This suggested that monomeric FANCI can displace one of the FANCD2 protomers in the homodimer. To test this directly, we immobilized FANCD2 on Streptavidin resin and incubated it with untagged FANCI. After washing, FANCI was bound to the beads in approximately 1:1 stoichiometric ratio with FANCD2 (Extended Data Fig. 6b). This is consistent with FANCI displacing one of the FANCD2 protomers to form the D2-I complex. Our results suggest a mechanism of autoinhibition wherein FANCD2 cannot be prematurely recruited to DNA prior to assembly with FANCI. This also raises the interesting possibility that the FANCD2-FANCI interaction may be regulated, and complex assembly is stimulated after DNA damage. Alternatively, it may be indicative of FANCI-independent functions of FANCD2. Interestingly, immunoprecipitation of FANCI from cells pulls down only ∼20% of FANCD2, suggesting that there may be a substantial pool of FANCD2 not bound to FANCI^9^.

In summary, our structural and biochemical analysis of FANCD2 and FANCI informs how DNA crosslink repair might be initiated (Fig. 4). FANCD2 adopts a closed conformation that does not interact with DNA. Upon incubation with FANCI, the FANCD2 homodimer readily exchanges into a stable D2-I heterodimer, which adopts an open conformation. D2-I binds DNA, and this causes a hinge-like rotation of FANCD2 and FANCI so they encircle double-stranded DNA (Fig. 4). Transition into this closed conformation exposes a region on the back of the FANCD2 hinge, allowing the FA core complex and UBE2T to access the monoubiquitination site. Ubiquitin acts as a covalent molecular pin to lock the ubD2-I DNA clamp into a closed conformation. Finally, the new FANCD2 surface exposed by DNA binding and the ubiquitinated lysine may act as a platform to recruit downstream DNA repair proteins, including the nuclease incision complex that removes the DNA lesion^6,11,12,27^. To release ubD2-I from DNA, the ubiquitin lock must be removed, providing an explanation for why the deubiquitination step is essential to complete repair^20^. This framework, informed by our structures of FANCD2 and FANCI, will be the focus of future work to fully understand the molecular basis of DNA interstrand crosslink repair.

**Fig. 4:**
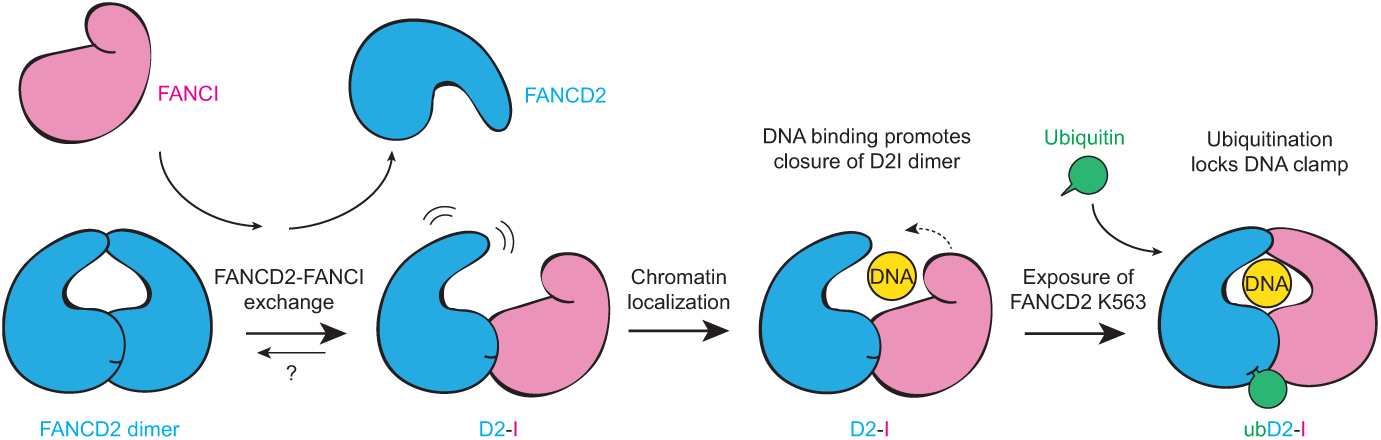
Model for regulation of FANCD2 and FANCI in DNA crosslink repair. FANCD2 purifies as a homodimer that is closed, does not bind DNA, and is not monoubiquitinated. Upon incubation with purified (monomeric) FANCI, this exchanges into a FANCD2-FANCI complex (D2-I) with an open conformation. D2-I binds and encircles DNA, converting the complex into a closed conformation, and thereby acting as a DNA clamp. The ubiquitination site on FANCD2 is exposed in the closed conformation, allowing access to the FA core complex and E2 enzyme. Ubiquitin locks the D2-I clamp in a closed conformation so it is not readily released from DNA.

## METHODS

### Cloning, expression and purification

cDNAs encoding full length *Gallus gallus* FANCI and FANCD2 were synthesized (GeneArt) and cloned into **pACEBac1**. The individual genes were amplified by PCR and cloned into **pBIG1a** vector using a modified version of the biGBac system, as previously described^28,29^. The combined vector was transformed into EMBacY *E. coli* competent cells for bacmid generation. The purified bacmid was then transfected into Sf9 cells. The virus was passaged twice before a large-scale culture was infected (5 ml of P2 virus into 500 ml of Sf9 cells at 2 million cells/ml). Cell growth and viability were monitored and cells harvested upon growth arrest (typically on day 3 after infection). A similar strategy was used for individual expression of FANCD2 and FANCI except that the *FANCD2* gene contained a C-terminal extension with a 3C protease site and double StrepII tag whereas the *FANCI* gene contained a C-terminal extension with a 3C protease site and 6x His tag.

For D2-I we took advantage of an excellent protein overexpression to devise a tag-free protein purification protocol based on sequential fractionation combining cation exchange and affinity chromatography. Cells were lysed by sonication in lysis buffer (100 mM HEPES pH 7.5, 300 mM NaCl, 1 mM TCEP, 5% glycerol, EDTA-free protease inhibitor, 5 mM benzamidine hydrochloride and 100 U/ml benzonase). Clarified cell lysate was passed through a HiTrap SP HP cation exchange chromatography column (GE Healthcare Life Sciences) to remove impurities. Flow through containing the unbound ID2 complex was diluted to 150 mM NaCl concentration and loaded onto a HiTrap Heparin HP affinity column (GE Healthcare Life Sciences). Using a shallow NaCl gradient, the ID2 complex was eluted around ∼ 500 mM NaCl concentration. The complex was run on a Superdex 200 26/60 column (GE Healthcare Life Sciences) in 50 mM HEPES pH 7.5, 150 mM NaCl and 1 mM TCEP. The fractions containing the complex were pooled and concentrated to ∼10 mg/ml and flash frozen for storage at −80 °C. Samples from each step were analyzed by SDS-PAGE using 4-12% NuPAGE Bis-Tris gels (ThermoFisher Scientific) to monitor the purification.

His-tagged FANCI was purified by sequential affinity and size exclusion chromatography. Clarified cell lysate produced as for D2-I was loaded onto a HisTrap HP column (GE Healthcare Life Sciences). Using an imidazole gradient, FANCI was eluted around ∼200 mM imidazole concentration. Collected fractions containing FANCI were diluted to ∼100 mM NaCl concentration and loaded onto a HiTrap Heparin HP affinity column (GE Healthcare Life Sciences). Using a shallow NaCl gradient, FANCI was eluted around ∼500 mM NaCl concentration. FANCI was then run on a Superdex 200 26/60 column (GE Healthcare Life Sciences) in 50 mM HEPES pH 7.5, 150 mM NaCl and 1 mM TCEP. The fractions containing FANCI were pooled and concentrated to ∼10 mg/ml and flash frozen for storage at −80 °C.

StrepII-tagged FANCD2 was also purified by sequential affinity and size exclusion chromatography. Clarified cell lysate produced as for D2-I and FANCI was incubated with Strep-Tactin Sepharose High Performance resin (GE Healthcare Life Sciences) for 60 min. The loaded resin was mounted on a glass column and washed twice with lysis buffer before elution with 8 mM D-Desthiobiotin. The elution was then diluted to ∼100 mM NaCl concentration and loaded onto a HiTrap Heparin HP affinity column (GE Healthcare Life Sciences). Using a shallow NaCl gradient, FANCD2 was eluted around ∼500 mM NaCl concentration. FANCD2 was then run on a Superdex 200 26/60 column (GE Healthcare Life Sciences) in 50 mM HEPES pH 7.5, 150 mM NaCl and 1 mM TCEP. The fractions containing FANCD2 were pooled and concentrated to ∼10 mg/ml and flash frozen for storage at −80 °C.

### *In vitro* ubiquitination and purification of ubiquitinated D2-I

Based on previously described ubiquitination assays^18,19^, we set up a large-scale *in vitro* reconstitution of the FA core complex-mediated ubiquitination of D2-I. In a total volume of 400 µl, the reaction contained 75 nM of E1 ubiquitin activating enzyme (Boston Biochem), 0.8 µM E2 (UBE2T)^18^, 0.5 µM E3 (FA core complex)^23^, 1 µM D2-I, 5 µM DNA (double stalled fork generated by annealing oligonucleotides X1, X2, X3, X4, X5, X6. Extended Data Table 2), 20 µM His-tagged ubiquitin (Enzo Life Sciences) in a reaction buffer of 50 mM HEPES pH 7.5, 64 mM NaCl, 4% glycerol, 5 mM MgCl_2_, 2 mM ATP and 0.5 mM DTT. The reaction was incubated at 30 °C for 90 min before applying it to 50 µl of Ni-NTA agarose resin (Qiagen) pre-equilibrated in W25 buffer (20 mM HEPES pH 7.5, 150 mM NaCl, 1 mM TCEP and 25 mM imidazole) in a 1.5 ml centrifuge tube. The resin was incubated under constant rotation at 4 °C for 60 min. The resin was washed twice, first with 400 µl of W25 buffer followed by a wash with 400 µl of W50 buffer (20 mM HEPES pH 7.5, 150 mM NaCl, 1 mM TCEP and 50 mM imidazole). Each wash was performed for 60 min at 4 °C under rotation. The Ni-NTA bound ubD2-I complex was eluted with W100 buffer (20 mM HEPES pH 7.5, 150 mM NaCl, 1 mM TCEP and 100 mM imidazole). Samples from each step were analyzed by SDS-PAGE to monitor the purification of ubD2-I complex. Presence of ubiquitin was confirmed by Western blot using anti-Ubiquitin (Millipore; Cat # 07-375). W100 elution fractions were then concentrated using a Vivaspin column (30 kDa MWCO) to a final volume of ∼20 µl and ∼500 nM concentration.

### Electrophoretic mobility shift assays (EMSAs)

Fluorescently labeled DNA (6FAM labeled on 3′ end, purchased from IDT) was prepared by incubating complementary oligonucleotides (Extended Data Table 2) at 95 °C for 5 min and slowly cooling down to room temperature over ∼2 h. For EMSAs, a 20 µl reaction containing 20 nM of 6FAM-DNA was incubated with the indicated concentration of protein in the presence of 50 mM HEPES pH 8.0, 150 mM NaCl and 1 mM TCEP. The reactions were incubated for 30 min at 22 °C. After incubation, 5 µl were directly loaded on a native polyacrylamide gel (6% DNA Retardation, Thermo Fisher) and run at 4 °C using 0.5 X TBE buffer for 60 min. The gel was then visualized using a Typhoon Imaging System (GE Healthcare). Each binding experiment was repeated three times and the intensity of free DNA was calculated using *ImageJ*^30^ for the quantification of the mean intensity and standard deviation between the measurements.

### Gel filtration assays

A volume of 300 µl of purified D2-I, FANCD2 and FANCI at 1 µM were sequentially run on a Superdex 200 10/300 column (GE Healthcare Life Sciences) equilibrated in 50 mM HEPES pH 7.5, 150 mM NaCl and 1 mM TCEP. To investigate the FANCD2-FANCI exchange we mixed and incubated 0.5 µM FANCD2 with 1 µM FANCI and FANCD2 for 30 min at room temperature in a total volume of 300 µl. The mix was then run on a Superdex 200 10/300 column (GE Healthcare Life Sciences) equilibrated in 50 mM HEPES pH 7.5, 150 mM NaCl and 1 mM TCEP. Fractions were analysed by SDS-PAGE using 4-12% NuPAGE Bis-Tris gels (ThermoFisher Scientific).

### Binding assays

We used purified StrepII-tagged FANCD2 and His-tagged FANCI to probe their interaction. In a total volume of 50 µl per reaction, FANCD2:FANCI at molar ratios 1:0, 1:0.5, 1:1, 1:2, 1:5 and 0:5 were mixed in 50 mM HEPES pH 7.5, 150 mM NaCl and 1 mM TCEP and incubated for 15 min at room temperature. Each reaction was applied to 20 µl of StrepTactin Sepharose High Performance resin (GE Healthcare Life Sciences) equilibrated in the same buffer. The loaded resin was then incubated for 30 min at 4 °C. The unbound fraction was removed and the resin further washed twice using 250 µl of the same buffer. The bound fraction was analyzed by SDS-PAGE using 4-12% NuPAGE Bis-Tris gels (ThermoFisher Scientific).

### Electron microscopy and image processing

Three microliters of ∼1 µM ubD2-I, D2-I or D2 complex were blotted on plasma cleaned UltraAufoil R1.2/1.3 grids^31^ (Quantifoil) for 3–4.5 s and plunged into liquid ethane using a Vitrobot Mark IV. The grids were imaged on a Titan Krios using Gatan K3 detector in super-resolution mode for ubD2-I and D2 or in counting mode for D2-I. Additional data was collected for ubD2-I at a tilt of 23° to overcome preferred orientation.

All image processing was performed using *Relion3*.*0/3*.*1*^32^ unless otherwise stated. The images were drift corrected using *MotionCorr2*^33^ and defocus was estimated using *CTFFIND4*^*34*^. Particles were initially picked manually and 2D classified. Selected classes from the 2D classification were used to autopick particles from the full datasets. After 2–3 rounds of 2D classification, classes with different orientations were selected for initial model generation in Relion. The initial model was used as reference for 3D classification into 4 classes. The selected classes from 3D classification were subjected to auto-refinement with solvent flattening using a soft mask. The defocus values were further refined using CTF Refinement in *Relion* followed by Bayesian polishing. Another round of auto-refinement was performed on these polished particles. All maps were post-processed to correct for modulation transfer function of the detector and sharpened by applying a negative B factor as determined automatically by *Relion*. A soft mask was applied during post processing to generate FSC curves to yield maps of average resolutions of 3.8 Å for ubD2-I, 4.1 Å for D2-I and 4.0 Å for D2.

To further improve the maps, we used focused classification and refinement of the ubD2-I and D2-I maps by dividing them into FANCI and FANCD2 regions. The density for DNA in ubD2-I was improved by using focused classification without image alignment.

To analyze the conformational heterogeneity in ubD2-I, we applied multi-body refinement using masks around FANCI and FANCD2 protomers. Three major motions were detected using principal component analysis on the optimal orientations of all the bodies for all particle images in the dataset using relion_flex_analyse^35^ (Supplementary Video 1).

### Modelling

The crystal structure of *Mus musculus* FANDC2-FANCI (PDB: 3S4W)^4^ was used as a template to generate a homology model of *Gallus gallus* D2-I in *I-TASSER*^36^. The homology model was initially fitted manually by visual inspection of the map followed by rigid fitting in *UCSF Chimera*^37^. The side chains were stubbed except for K563 of FANCD2 and the model was iteratively refined in *Coot*^38,39^ and *Phenix*^40^. The crystal structure of ubiquitin (PDB: 1UBQ)^41^ was rigidly fitted into the unassigned density in the ubD2-I map and refined in *Coot*. An idealized dsDNA of 33 bp length was placed and refined in the density observed for DNA in ubD2-I using Coot.

### Data availability

CryoEM maps generated during this study have been deposited in the Electron Microscopy Data Bank (EMDB) with accession codes EMD-XXXX (D2-I), EMD-YYYY (ubD2-I), and EMD-ZZZZ (D2-D2). Models generated during this study have been deposited in the protein databank (PDB) with accession codes #### (D2-I) and #### (ubD2-I). All other data are available from the authors upon reasonable request.

## Acknowledgements

We are grateful to Takanore Nakane, Sjors Scheres, Madan Babu, Paul Emsley and members of the Passmore lab for assistance and advice; the LMB EM facility, Jake Grimmett and Toby Darling (LMB scientific computation), and Jianguo Shi (baculovirus) for support. This work was supported by the Medical Research Council, as part of United Kingdom Research and Innovation, MRC file reference number MC_U105192715 (L.A.P). P.A. is supported by an EMBO Long Term Fellowship (ALTF 692-2018). We acknowledge Diamond Light Source for access to eBIC (proposals ####) funded by the Wellcome Trust, MRC, and Biotechnology and Biological Sciences Research Council.

## Author contributions

P.A. and S.S. designed protein expression and purification schemes, performed ubiquitination and binding assays, performed cryoEM, 3D reconstruction and modelling; L.A.P. and K.J.P. supervised the research; all authors contributed to writing the paper.

## Author information

The authors declare no competing interests. Readers are welcome to comment on the online version of the paper. Correspondence and requests for materials should be addressed to.

## EXTENDED DATA

**Extended Data Table 1.** Cryo-EM data collection and refinement.

|  | D2-I<br>EMD-XXXXX,<br>PDB XXXX | ubD2-I<br>EMD-XXXXX<br>PDB XXXX | D2<br>EMD-XXXXX |
| --- | --- | --- | --- |
| **Data collection and processing** |  |  |  |
| Magnification | 81,000 X | 81,000 X | 81,000 X |
| Voltage (keV) | 300 | 300 | 300 |
| Electron exposure (e-/Å<sup>2</sup>) | ~70 | ~70 | ~70 |
| Defocus range (μm) | -1.8 to -3.5 | -1.8 to -3.5 | -1.8 to -3.5 |
| Pixel size (Å) | 1.11 (LMB) | 1.06 (eBIC), 1.11 (LMB) | 1.11 (eBIC) |
| Symmetry imposed | C1 | C1 | C1 |
| Initial particle images (no.) | ~2,000,000 | ~2,500,000 | ~3,000,000 |
| Final particle images (no.) | 171,936 | 146,245 | 901,085 |
| Map resolution (Å) | 4.1 | 3.8 | 4.0 |
| FSC threshold | 0.143 | 0.143 | 0.143 |
| Map resolution range (Å) | 4.1 to > 10 | 3.8 to > 10 | 4.0 to > 10 |

**Extended Data Table 2.**
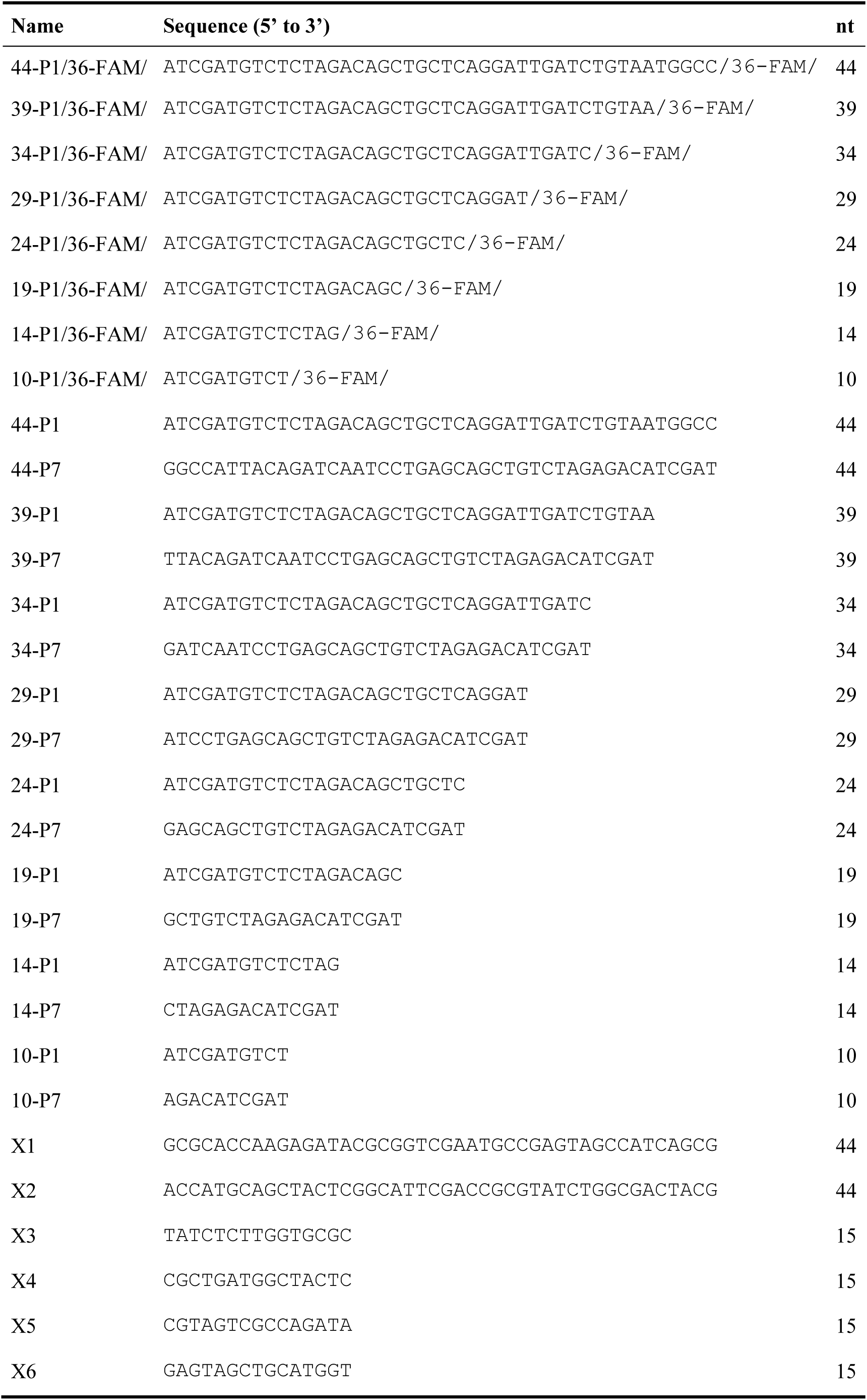
DNA oligonucleotides used in this study.

**Extended Data Fig. 1:**
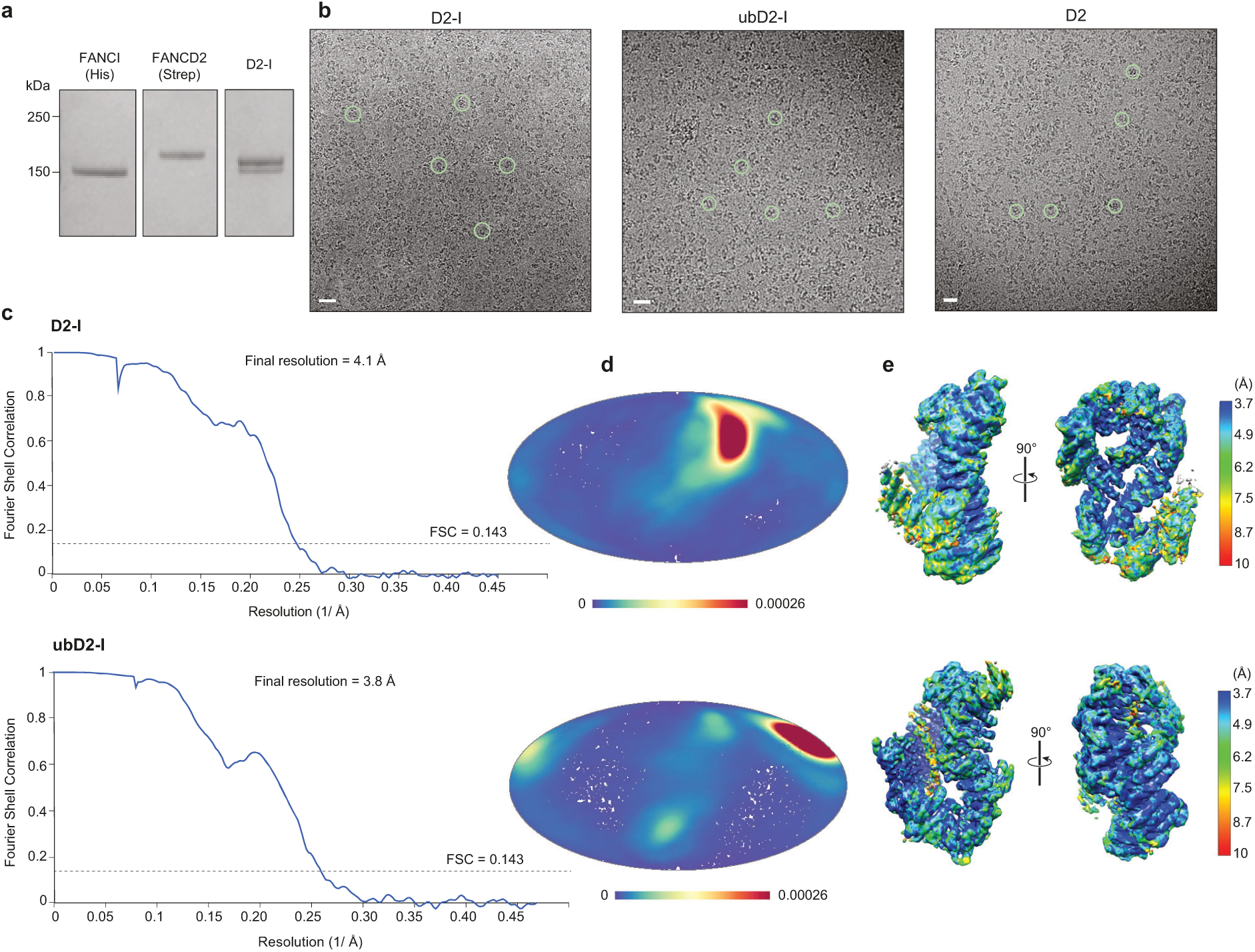
Purification of FANCI, FANCD2, and D2-I, and cryoEM of D2-I, ubD2-I and D2 dimer. **a** Coomassie gel showing purified His-tagged FANCI, StrepII-tagged FANCD2 and D2-I after gel filtration. **b-e** CryoEM of D2-I, ubD2-I and D2. **b** Representative micrographs. Selected individual particles are marked with green circles. Scale bar is 25 nm. **c** Fourier shell correlation curves for gold-standard refinements. **d** Angular distribution density plots of particles used in 3D reconstructions calculated using *cryoEF*^42^. Every point is a particle orientation and the color scale represents the normalized density of views around this point. The color scale runs from 0 (low, blue) to 0.00026 (high, red). All complexes had a preferred orientation. Note that C2 symmetry was applied for FANCD2. **e** Local resolution estimates calculated using *ResMap*^43^.

**Extended Data Fig. 2.**
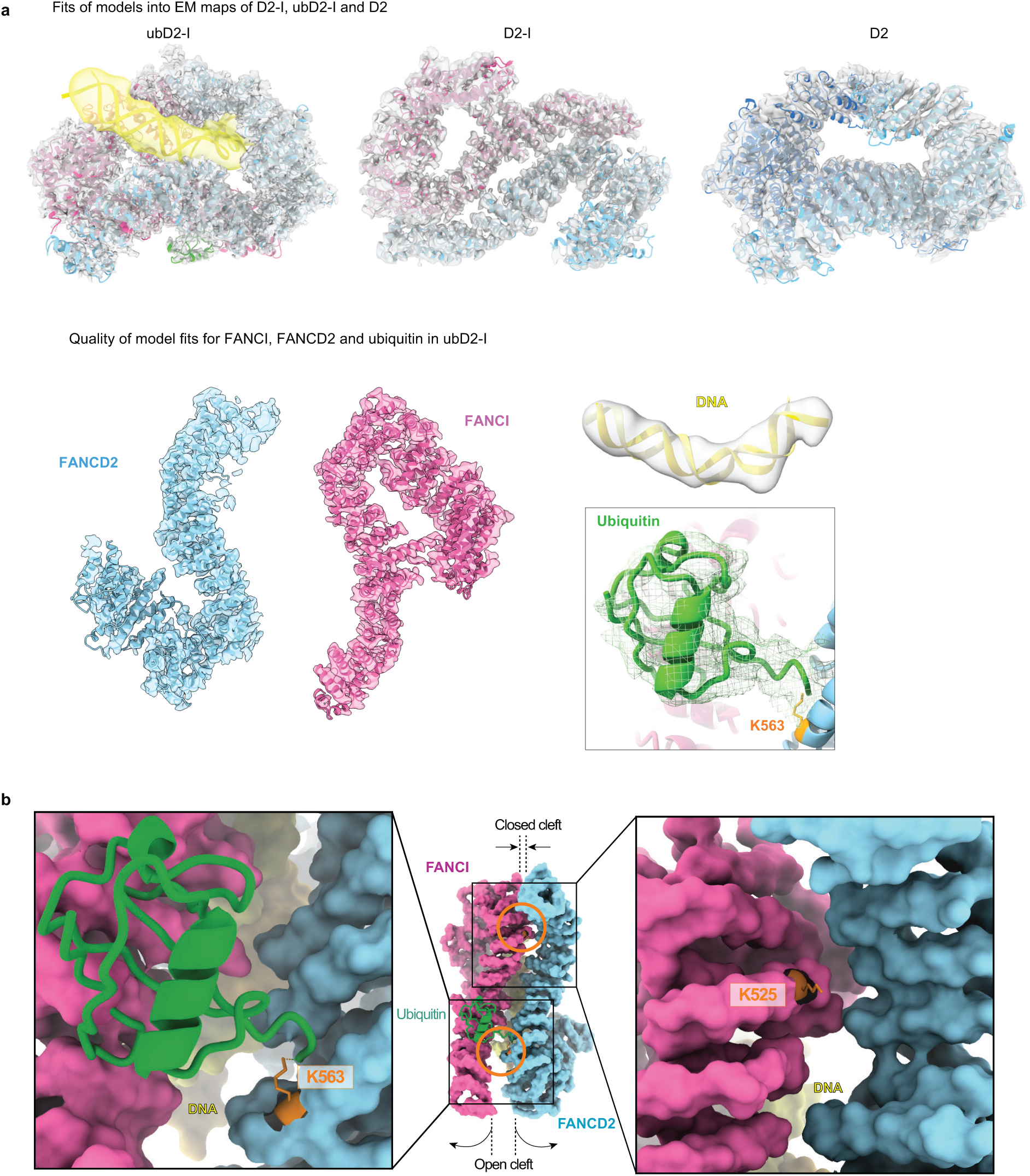
Model fitting. **a** Representative fits of model to map. **b** Details of monoubiquitinated lysines (orange) in FANCD2 (blue) and FANCI (magenta) within the ubD2-I structure, shown as a surface representation of the model for FANCD2 and FANCI, and in cartoon for ubiquitin.

**Extended Data Fig. 3.**
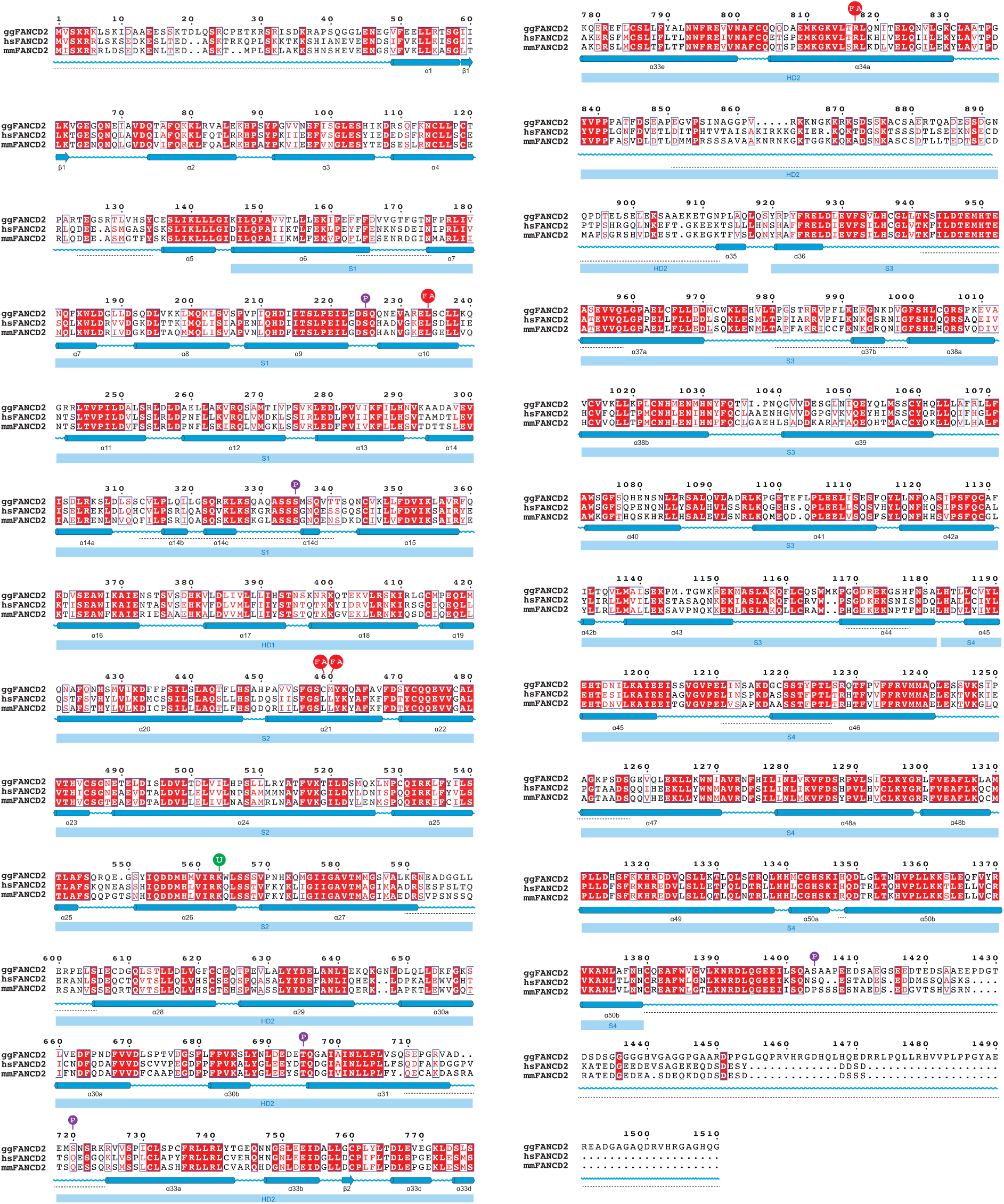
Sequence alignment of FANCD2. Sequences from *Gallus gallus* (gg), *Homo sapiens* (hs) and *Mus musculus* (mm) are shown. Under the sequence alignment, the secondary structure of ggFANCD2 is represented in blue: tubes for α-helices and arrows for β-strands. For comparison, the helices have been labelled according to their equivalent nomenclature in the murine structure of unmodified D2-I, and the solenoid (S) and helical domain (HD) numbers are indicated in light blue boxes^4^. Red signs (“FA”) indicate mutations characterized in FA patients, purple signs (“P”) indicate phosphorylation sites and the monoubiquitination site K563 is marked in green (“U”). Disordered regions are marked with dashed lines.

**Extended Data Fig. 4.**
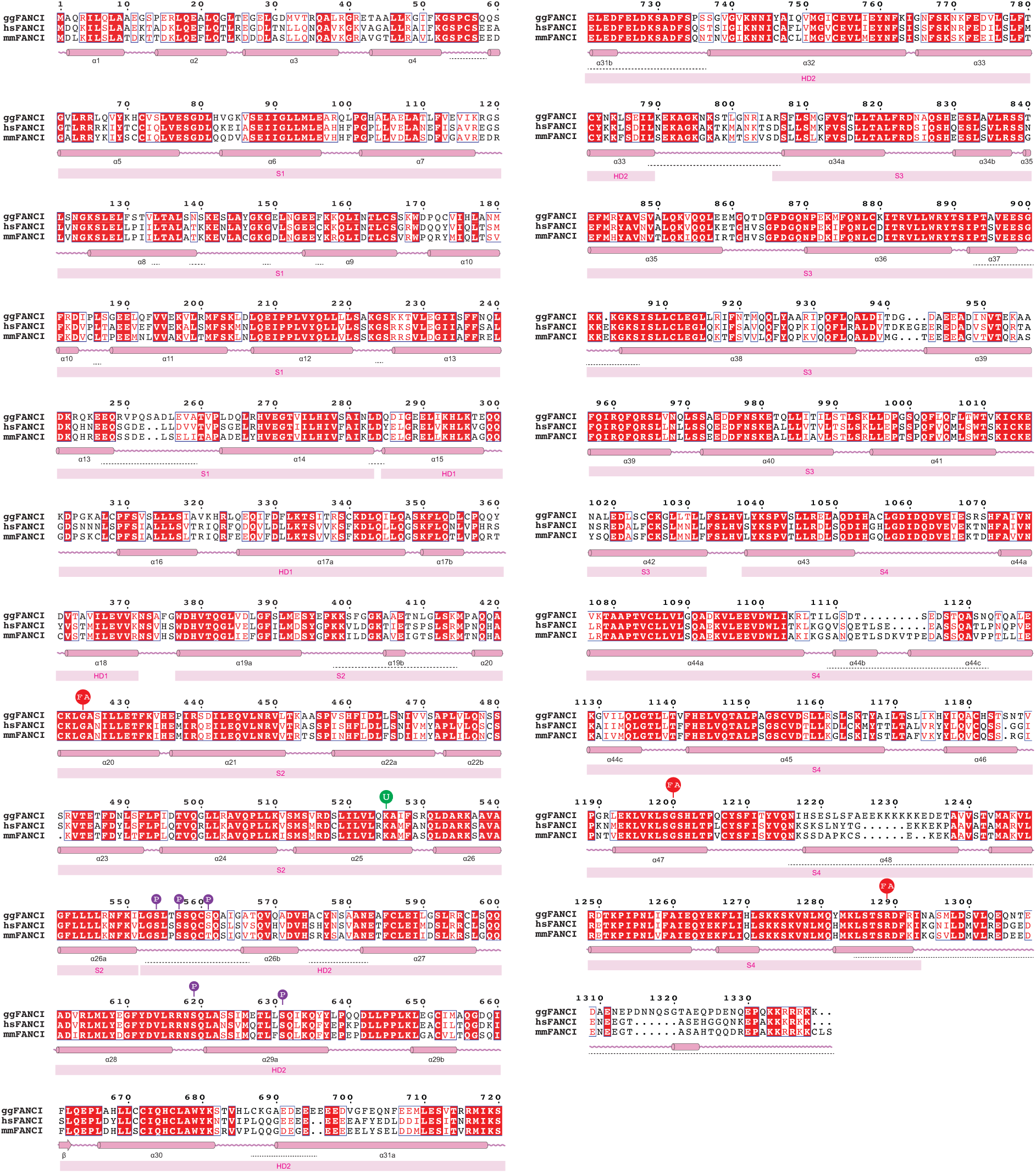
Sequence alignment of FANCI. Sequences from *Gallus gallus* (gg), *Homo sapiens* (hs) and *Mus musculus* (mm) are shown. Under the sequence alignment, the secondary structure of ggFANCI is represented in blue: tubes for α-helices and arrows for β-strands. For comparison, the helices have been labelled according to their equivalent nomenclature in the murine structure of unmodified D2-I, and the solenoid (S) and helical domain (HD) numbers are indicated in light blue boxes^4^. Red signs (“FA”) indicate mutations characterized in FA patients, purple signs (“P”) indicate phosphorylation sites and the monoubiquitination site K525 is marked in green (“U”). Disordered regions are marked with dashed lines.

**Extended Data Fig. 5.**
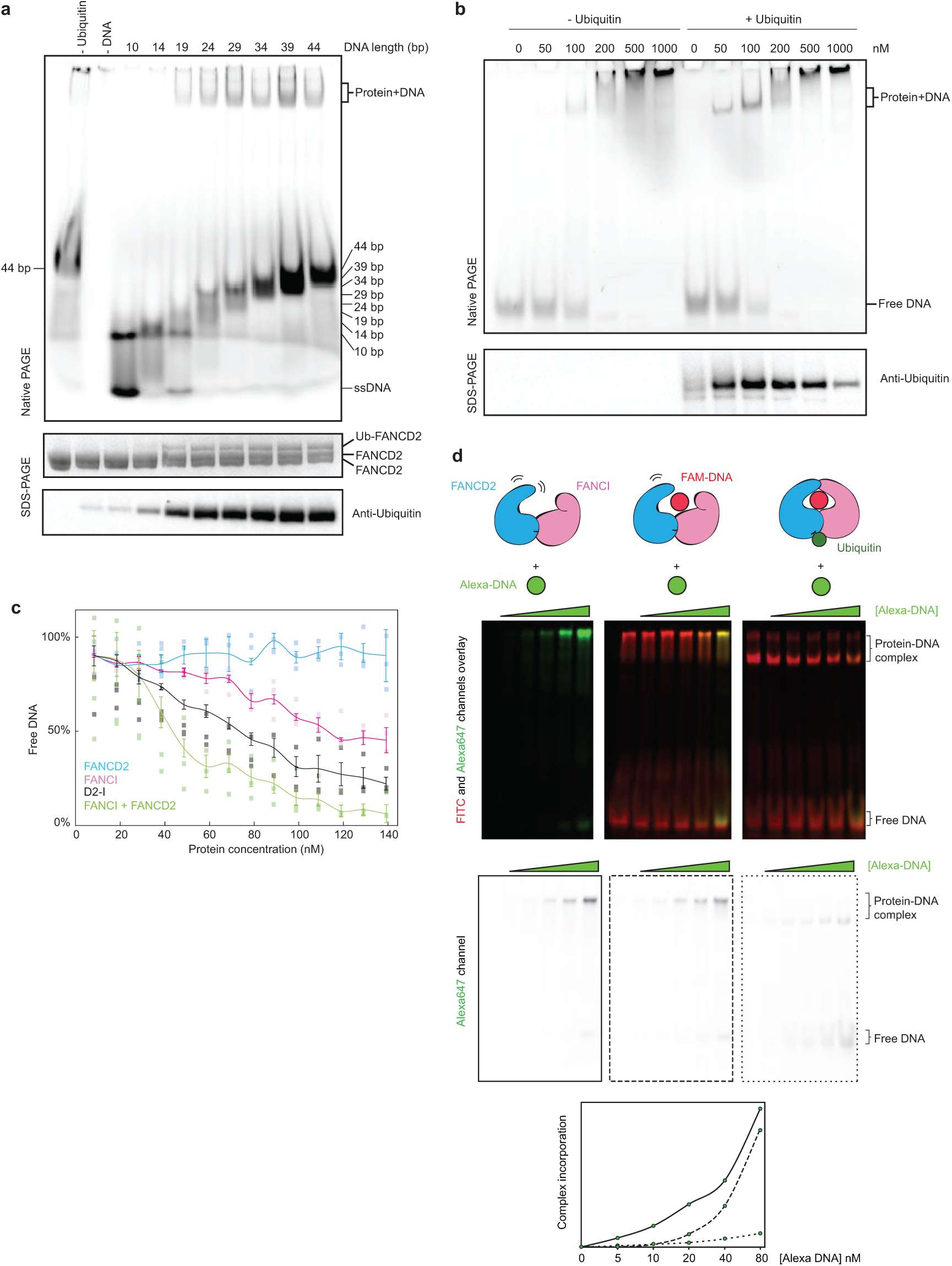
Analysis of DNA binding by FANCD2 and FANCI. **a** Monoubiquitination assays were assembled in the presence of linear double-stranded DNA of differing lengths (10–44 bp). DNA binding was analyzed by EMSA (top) and imaging of the fluorescently labeled DNA. Monoubiquitination efficiency was analyzed by Coomassie blue (middle) and Western blotting the His-tagged ubiquitin (bottom). Controls lacking ubiquitin or DNA are indicated. **b** Monoubiquitination assays of D2-I were assembled without (left) and with (right) ubiquitin, both in the presence of increasing amounts of a 39 bp double-stranded DNA (0 – 1000 nM). Assays were analyzed by EMSA (top, imaging for the fluorescently-labeled DNA) or Western blotting His-tagged ubiquitin (bottom). These data are representative of experiments performed twice. **c** DNA binding of FANCI, FANCD2, D2-I, and FANCI mixed with FANCD2 was analyzed by EMSAs performed with 20 nM 39 bp double-stranded DNA and 0–140 nM protein. Representative gels of experiments independently performed three times (Fig. 3b) and quantitation of mean intensity of free DNA are shown. Error bars represent the standard deviation. Individual data points are shown and means connected by lines for clarity.

**Extended Data Fig. 6.**
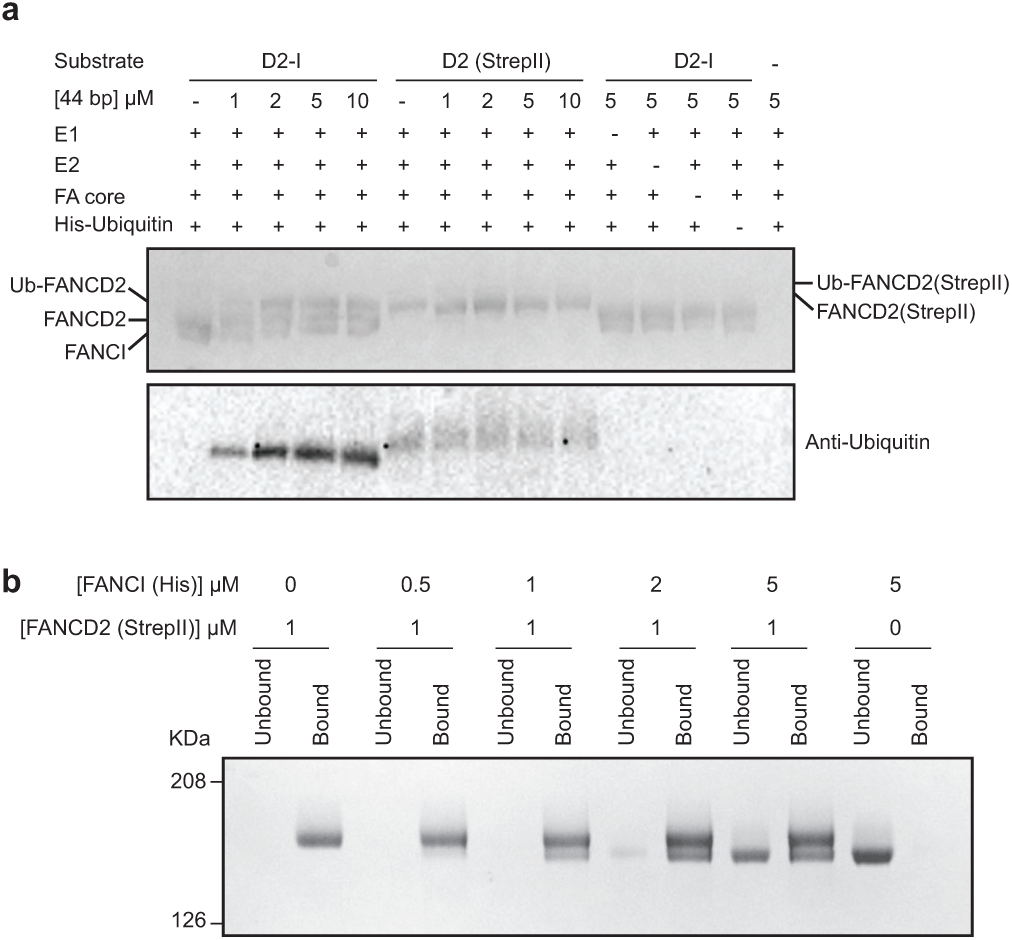
FANCD2 forms a homodimer that does not bind DNA. **a** Monoubiquitination assays of D2-I and FANCD2 homodimer. The FANCD2 homodimer had a StrepII-tag and was therefore larger than D2 in the D2-I complex. Monoubiquitination efficiency was analyzed by Coomassie blue SDS-PAGE (top) and Western blotting the His-tagged ubiquitin (bottom). **b** FANCD2/FANCI exchange assay. The FANCD2 homodimer was immobilized on Streptactin resin and incubated with free FANCI. The resin was washed, then bound and unbound fractions were analyzed by SDS-PAGE. These data are representative of experiments performed twice.

## REFERENCES

1 Kottemann, M. C. & Smogorzewska, A. Fanconi anaemia and the repair of Watson and Crick DNA crosslinks. Nature 493, 356–363, doi:10.1038/nature11863 (2013).

2 Crossan, G. P. & Patel, K. J. The Fanconi anaemia pathway orchestrates incisions at sites of crosslinked DNA. J Pathol 226, 326–337, doi:10.1002/path.3002 (2012).

3 Walden, H. & Deans, A. J. The Fanconi anemia DNA repair pathway: structural and functional insights into a complex disorder. Annu Rev Biophys 43, 257–278, doi:10.1146/annurev-biophys-051013-022737 (2014).

4 Joo, W. et al. Structure of the FANCI-FANCD2 complex: insights into the Fanconi anemia DNA repair pathway. Science 333, 312–316, doi:10.1126/science.1205805 (2011).

5 Garcia-Higuera, I. et al. Interaction of the fanconi anemia proteins and BRCA1 in a common pathway. Molecular Cell 7, 249–262, doi:Doi 10.1016/S1097-2765(01)00173-3 (2001).

6 Knipscheer, P. et al. The Fanconi anemia pathway promotes replication-dependent DNA interstrand cross-link repair. Science 326, 1698–1701, doi:10.1126/science.1182372 (2009).

7 Meetei, A. R. et al. A multiprotein nuclear complex connects Fanconi anemia and Bloom syndrome. Mol Cell Biol 23, 3417–3426, doi:10.1128/mcb.23.10.3417-3426.2003 (2003).

8 Sims, A. E. et al. FANCI is a second monoubiquitinated member of the Fanconi anemia pathway. Nat Struct Mol Biol 14, 564–567, doi:10.1038/nsmb1252 (2007).

9 Smogorzewska, A. et al. Identification of the FANCI protein, a monoubiquitinated FANCD2 paralog required for DNA repair. Cell 129, 289–301, doi:10.1016/j.cell.2007.03.009 (2007).

10 Montes de Oca, R. et al. Regulated interaction of the Fanconi anemia protein, FANCD2, with chromatin. Blood 105, 1003–1009, doi:10.1182/blood-2003-11-3997 (2005).

11 MacKay, C. et al. Identification of KIAA1018/FAN1, a DNA repair nuclease recruited to DNA damage by monoubiquitinated FANCD2. Cell 142, 65–76, doi:10.1016/j.cell.2010.06.021 (2010).

12 Liu, T., Ghosal, G., Yuan, J., Chen, J. & Huang, J. FAN1 acts with FANCI-FANCD2 to promote DNA interstrand cross-link repair. Science 329, 693–696, doi:10.1126/science.1192656 (2010).

13 Smogorzewska, A. et al. A genetic screen identifies FAN1, a Fanconi anemia-associated nuclease necessary for DNA interstrand crosslink repair. Mol Cell 39, 36–47, doi:10.1016/j.molcel.2010.06.023 (2010).

14 Kratz, K. et al. Deficiency of FANCD2-associated nuclease KIAA1018/FAN1 sensitizes cells to interstrand crosslinking agents. Cell 142, 77–88, doi:10.1016/j.cell.2010.06.022 (2010).

15 Klein Douwel, D. et al. XPF-ERCC1 acts in Unhooking DNA interstrand crosslinks in cooperation with FANCD2 and FANCP/SLX4. Mol Cell 54, 460–471, doi:10.1016/j.molcel.2014.03.015 (2014).

16 Hodskinson, M. R. et al. Mouse SLX4 is a tumor suppressor that stimulates the activity of the nuclease XPF-ERCC1 in DNA crosslink repair. Mol Cell 54, 472–484, doi:10.1016/j.molcel.2014.03.014 (2014).

17 Yamamoto, K. N. et al. Involvement of SLX4 in interstrand cross-link repair is regulated by the Fanconi anemia pathway. Proc Natl Acad Sci U S A 108, 6492–6496, doi:10.1073/pnas.1018487108 (2011).

18 Rajendra, E. et al. The genetic and biochemical basis of FANCD2 monoubiquitination. Mol Cell 54, 858–869, doi:10.1016/j.molcel.2014.05.001 (2014).

19 Sato, K., Toda, K., Ishiai, M., Takata, M. & Kurumizaka, H. DNA robustly stimulates FANCD2 monoubiquitylation in the complex with FANCI. Nucleic Acids Res 40, 4553–4561, doi:10.1093/nar/gks053 (2012).

20 Oestergaard, V. H. et al. Deubiquitination of FANCD2 is required for DNA crosslink repair. Mol Cell 28, 798–809, doi:10.1016/j.molcel.2007.09.020 (2007).

21 Kim, J. M. et al. Inactivation of murine Usp1 results in genomic instability and a Fanconi anemia phenotype. Dev Cell 16, 314–320, doi:10.1016/j.devcel.2009.01.001 (2009).

22 Nijman, S. M. et al. The deubiquitinating enzyme USP1 regulates the Fanconi anemia pathway. Mol Cell 17, 331–339, doi:10.1016/j.molcel.2005.01.008 (2005).

23 Shakeel, S. et al. Structure of the Fanconi anemia monoubiquitin ligase complex. Nature 575, 234–237 (2019) doi:10.1038/s41586-019-1703-4.

24 Swuec, P. et al. The FA Core Complex Contains a Homo-dimeric Catalytic Module for the Symmetric Mono-ubiquitination of FANCI-FANCD2. Cell Rep 18, 611–623, doi:10.1016/j.celrep.2016.11.013 (2017).

25 van Twest, S. et al. Mechanism of Ubiquitination and Deubiquitination in the Fanconi Anemia Pathway. Mol Cell 65, 247–259, doi:10.1016/j.molcel.2016.11.005 (2017).

26 Ishiai, M. et al. FANCI phosphorylation functions as a molecular switch to turn on the Fanconi anemia pathway. Nat Struct Mol Biol 15, 1138–1146, doi:10.1038/nsmb.1504 (2008).

27 Crossan, G. P. et al. Disruption of mouse Slx4, a regulator of structure-specific nucleases, phenocopies Fanconi anemia. Nat Genet 43, 147–152, doi:10.1038/ng.752 (2011).

## ONLINE REFERENCES

28 Weissmann, F. et al. biGBac enables rapid gene assembly for the expression of large multisubunit protein complexes. Proc Natl Acad Sci U S A 113, E2564–2569, doi:10.1073/pnas.1604935113 (2016).

29 Hill, C. H. et al. Activation of the Endonuclease that Defines mRNA 3’ Ends Requires Incorporation into an 8-Subunit Core Cleavage and Polyadenylation Factor Complex. Mol Cell, doi:10.1016/j.molcel.2018.12.023 (2019).

30 Rueden, C. T. et al. ImageJ2: ImageJ for the next generation of scientific image data. BMC Bioinformatics 18, 529, doi:10.1186/s12859-017-1934-z (2017).

31 Russo, C. J. & Passmore, L. A. Electron microscopy: Ultrastable gold substrates for electron cryomicroscopy. Science 346, 1377–1380, doi:10.1126/science.1259530 (2014).

32 Zivanov, J. et al. New tools for automated high-resolution cryo-EM structure determination in RELION-3. Elife 7, doi:10.7554/eLife.42166 (2018).

33 Zimmerman, E. S., Schulman, B. A. & Zheng, N. Structural assembly of cullin-RING ubiquitin ligase complexes. Curr Opin Struct Biol 20, 714–721, doi:10.1016/j.sbi.2010.08.010 (2010).

34 Rohou, A. & Grigorieff, N. CTFFIND4: Fast and accurate defocus estimation from electron micrographs. J Struct Biol 192, 216–221, doi:10.1016/j.jsb.2015.08.008 (2015).

35 Nakane, T., Kimanius, D., Lindahl, E. & Scheres, S. H. Characterisation of molecular motions in cryo-EM single-particle data by multi-body refinement in RELION. Elife 7, doi:10.7554/eLife.36861 (2018).

36 Yang, J. et al. The I-TASSER Suite: protein structure and function prediction. Nat Methods 12, 7–8, doi:10.1038/nmeth.3213 (2015).

37 Pettersen, E. F. et al. UCSF Chimera--a visualization system for exploratory research and analysis. J Comput Chem 25, 1605–1612, doi:10.1002/jcc.20084 (2004).

38 Emsley, P., Lohkamp, B., Scott, W. G. & Cowtan, K. Features and development of Coot. Acta Crystallogr D Biol Crystallogr 66, 486–501, doi:10.1107/S0907444910007493 (2010).

39 Emsley, P. & Cowtan, K. Coot: model-building tools for molecular graphics. Acta Crystallogr D Biol Crystallogr 60, 2126–2132, doi:10.1107/S0907444904019158 (2004).

40 Adams, P. D. et al. PHENIX: a comprehensive Python-based system for macromolecular structure solution. Acta Crystallogr D Biol Crystallogr 66, 213–221, doi:10.1107/S0907444909052925 (2010).

41 Vijay-Kumar, S., Bugg, C. E. & Cook, W. J. Structure of ubiquitin refined at 1.8 A resolution. J Mol Biol 194, 531–544, doi:10.1016/0022-2836(87)90679-6 (1987).

42 Naydenova, K. & Russo, C. J. Measuring the effects of particle orientation to improve the efficiency of electron cryomicroscopy. Nat Commun 8, 629, doi:10.1038/s41467-017-00782-3 (2017).

43 Kucukelbir, A., Sigworth, F. J. & Tagare, H. D. Quantifying the local resolution of cryo-EM density maps. Nat Methods 11, 63–65, doi:10.1038/nmeth.2727 (2014).

